# PARylation in Parkinson’s disease: a bridge between Lewy body formation and neuronal cell death

**DOI:** 10.1101/2025.03.12.642849

**Authors:** Claudia Novello, Federica Giampietro, Alessandra Maria Calogero, Giorgio Giaccone, Michele Salemi, Manuela Bramerio, Emanuela Bonoldi, Daniela Calandrella, Elena Contaldi, Ioannis Ugo Isaias, Chiara Rolando, Gianni Pezzoli, Graziella Cappelletti, Samanta Mazzetti

## Abstract

Poly-ADP-ribosylation (PARylation), catalyzed by the enzyme PARP1, involves the addition of poly-ADP-ribose polymers (PAR) and has been associated with α-synuclein aggregation in Parkinson’s disease (PD) models. This study aimed to unravel the role of PARylation in α-synuclein aggregation and neuronal cell death in the complex environment of *post-mortem* human PD brains. Using high-resolution imaging and 3D reconstruction analysis, we observed that PAR accumulate in the cytoplasm in regions affected by PD pathology, preceding the formation of α-synuclein oligomers. Additionally, we found that PAR and stress granules contribute to the formation of Lewy bodies. Increased colocalization of PAR with mitochondria in the substantia nigra of PD patients, along with the presence of PAR-positive condensed DNA, further suggests a role in neuronal cell death.

Collectively, our findings reveal a critical involvement of PARylation in the pathological mechanisms underlying neurodegeneration in PD and position PARylation as a potential therapeutic target.

**GRAPHICAL ABSTRACT:** 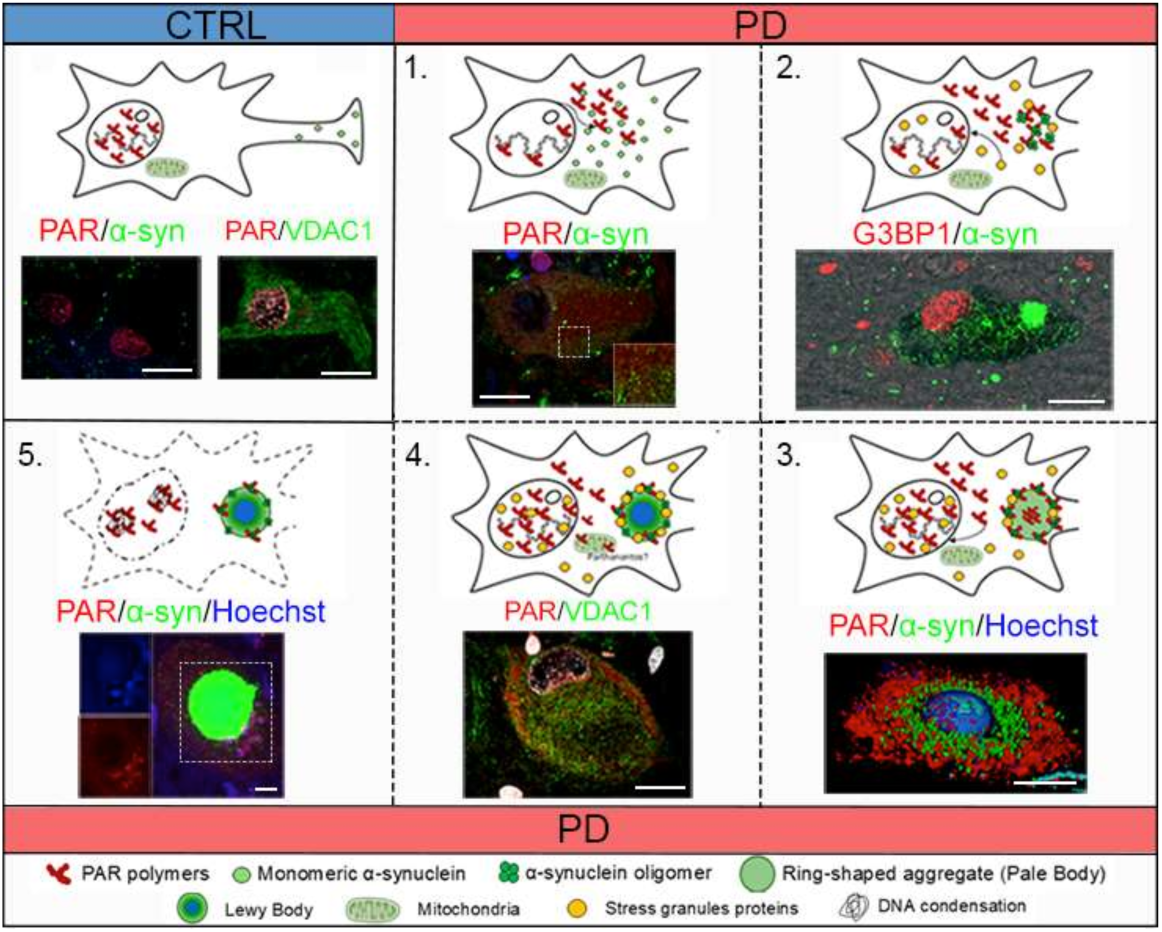

## 1 Introduction

Parkinson’s disease (PD) is the most common neurodegenerative movement disorder associated with the loss of *substantia nigra* dopaminergic neurons and the presence of intraneuronal inclusions known as Lewy bodies (LBs) and Lewy neurites^1,2^, which are enriched in α-synuclein protein^3^. Many aspects of PD pathogenesis remain unclear, particularly the mechanisms driving α-synuclein aggregation and neuronal death^4^.

Special attention is paid to the role of post-translational modifications (PTMs) in neurodegeneration, as they are crucial in influencing structure, function, and biochemical properties of proteins^5^. Interestingly, *in vitro* and in mouse model evidence revealed that pathological accumulation of α-synuclein could be driven by PARP1- mediated Poly-ADP-ribosylation (PARylation)^6^, a PTM consisting in the covalent addition of poly-ADP-ribose units to the carboxyl group of acidic residues on target proteins. Poly-ADP-ribose polymers (PAR) formation is catalysed by the nuclear enzyme PARP17,8, that consists of three domains: *i)* N-terminal DNA-binding domain, *ii)* central auto-modification domain, and *iii)* C-terminal catalytic domain (CAT)7,8. Physiologically, PARP1-mediated PARylation is fundamental for several cellular pathways such as protein stability^9–11^, gene expression^12^, inflammation^13^, and autophagy^14^, but it is particularly known for its dual function in regulating cellular response to oxidative stress^15,16^, which is reported to be elevated in PD^17^. In *in vitro* stressed cell models, cytoplasmic PAR increase determines the formation of stress granules^18^ through the PARylation of stress granules proteins, such as Ras-GTPase-activating protein (GAP)-binding protein 1 (G3BP1)^19^. While these findings are highly informative, there is a lack of evidence demonstrating PAR’s role in the human PD brain and how it regulates various stress-related cellular processes, including stress granule formation and cell death, *in vivo*.

PARP1 plays a pivotal role in regulating cell function and survival. It is essential for DNA damage repair^16,20^; but its overactivation in response to severe DNA damage can lead to Parthanatos pathway^21^. Parthanatos is a caspase-independent cell death mechanism characterized by PAR overproduction and accumulation into the cytoplasm. This process is linked to mitochondrial release of apoptosis-inducing factor (AIF), followed by the nuclear translocation of the AIF/macrophage migration inhibitory factor (MIF) complex, which leads to MIF-mediated DNA fragmentation^22,23^.

Emerging evidence links PARP1 overactivation to α-synuclein aggregation and Parthanatos pathway activation in a PD mouse model^6,24^. These data are supported by the observation of an increase in PAR levels, concomitantly with a α-synuclein decrease, in the cerebrospinal fluid of PD patients compared with control subjects^6,25,26^. In addition, in *post-mortem* human brain obtained from PD patients, PARP1 translocates from the nucleus to the cytoplasm of the *substantia nigra* dopaminergic neurons where it colocalizes with α-synuclein and accumulates into LBs^27^. Lastly, PAR were found to colocalize with phosphorylated α-synuclein^28^. These studies suggest a key role for PARylation of α-synuclein in neurodegeneration but its involvement in the early stages of aggregation is still unexplored, leading us to investigate this process in *post-mortem* human brain from PD patients. In the complexity of human brain, we aimed to disentangle: *i)* the interplay of PARylation and α-synuclein aggregation from the initial appearance of α-synuclein oligomers to mature LB; *ii)* the impact of PARylation in the response to cellular stresses, specifically the formation of stress granules; *iii)* the role of PARylation in neuronal cell death, focusing on PARylation of both mitochondria and condensed DNA, specifically involved in Parthanatos pathway. We found cytoplasmic accumulation of PAR in the brain regions mirroring previously reported PARP1 activity in PD human brain^27^. We also linked cytoplasmic PARylation with LB formation, including α-synuclein oligomers, which are the earliest aggregates^29^ providing crucial evidence that PARylation drives α-synuclein aggregation^6^ in the human brain. Furthermore, our analysis of G3BP1 and TIA1-related proteins (TIAR), which are markers of stress granule proteins, revealed their redistribution during LB formation^18^. Finally, we pointed out an increase of PARylation within mitochondria and the presence of condensed DNA, suggesting the activation of Parthanatos pathway that could be responsible for dopaminergic neurons cell death in PD human brain. All together these data added important pieces to understand how α-synuclein can be involved in neurodegeneration and contribute to the emerging view that PARylation could be a promising future therapeutic target.

## 2 Results

### 2.1 PAR neuronal redistribution is linked to α-synuclein aggregation in brain of PD patients

*In vitro* studies and genetic mouse model showed that PARP1 and PAR are involved in PD pathogenesis, including the conversion of α-synuclein to a more toxic species and neuron death^6,25,27^. In the present study we aimed to investigate whether PARylation, an index for PARP enzymatic activity, may be altered in human PD brain and how it can influence disease pathophysiology. First, we analysed whether PARP1 redistribution from the nucleus to the cytoplasm of neurons and astrocytes that we previously found in PD patients^27^ also affects the localization of PAR. For this purpose, we performed immunoenzymatic assay for PAR. In control subjects, PAR was predominantly localised in the nuclei of neurons (<u>Fig. 1a</u>, black arrowheads). However, in PD patients, we observed a redistribution of PAR from the nucleus to the cytoplasm, consistent with the previously reported pattern for PARP1 (<u>Fig. 1b</u>, black arrows). We quantified the number of PAR-positive nuclei and found a significant reduction in PD patients compared to controls in the frontal cortex, entorhinal cortex, and cingulate cortex (<u>Fig. 1c-e</u>). In contrast, no significant differences were found between controls and PD subjects in the percentage of PAR-positive nuclei in cortical astrocytes, where PAR remained predominantly localised in the nucleus with little to no presence in their cellular body (Supplementary Fig. 1c). We also investigated PAR redistribution in astrocytes and neurons in the *substantia nigra*. As for frontal cortex, we reported no differences in PAR positive nuclei of astrocytes between controls and PD subjects (Supplementary <u>Fig. 1a-b”, d</u>). Regarding neuronal PAR redistribution (<u>Fig.1 f, g</u>), while most PD patients exhibited a reduction in PAR-positive nuclei compared to controls, this decrease was not statistically significant (<u>Fig. 1h</u>).

**Fig. 1.**
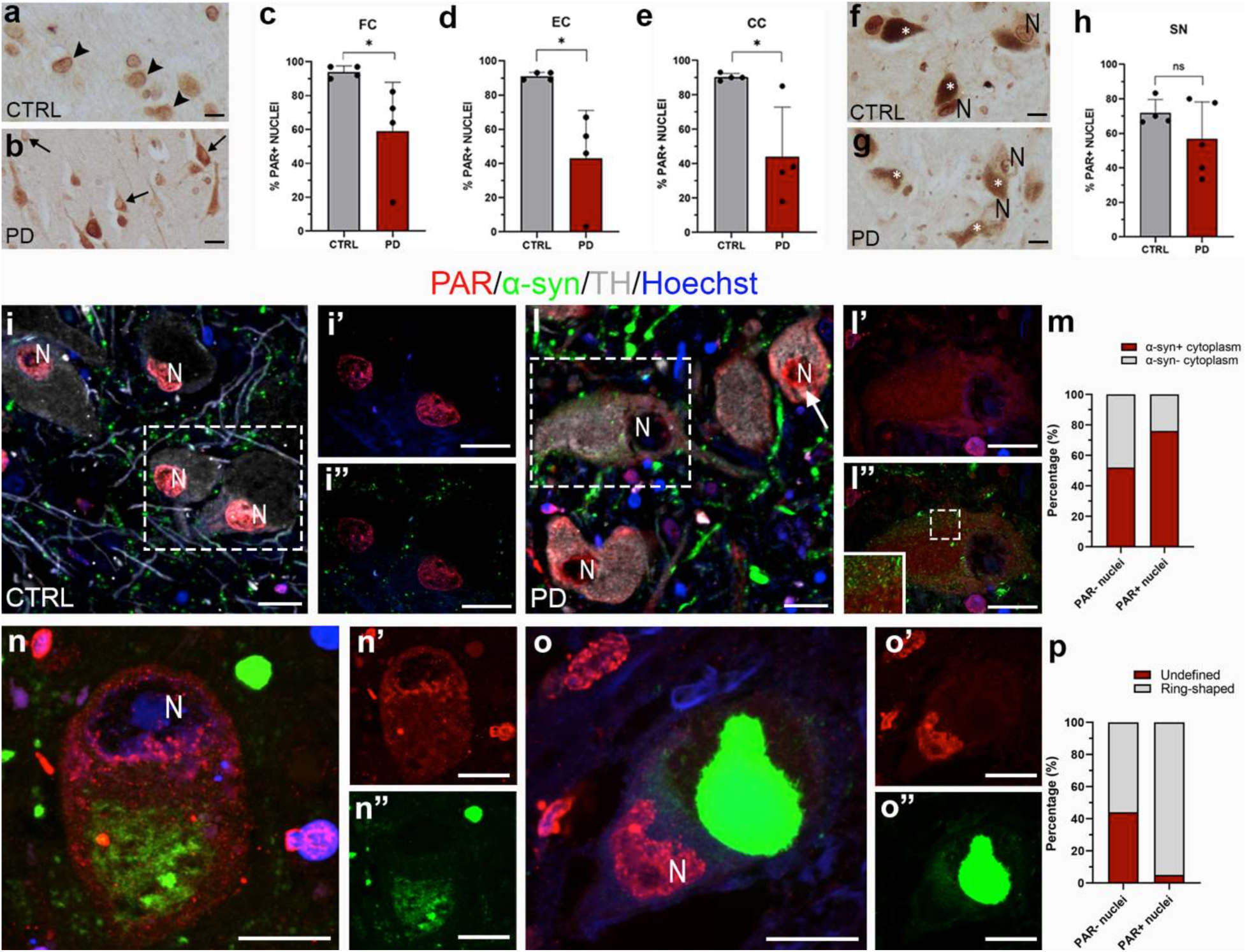
PAR distribution changes in *post-mortem* human brain in PD patients compared to control subjects. (**a, b**) representative image of PAR distribution in human frontal cortex. In control samples, PAR is localized in the nuclei of neurons (black arrowheads), while in PD patients a redistribution from nucleus to cytoplasm can be observed (black arrows). (**c, e**) the graphs show the percentage of PAR positive neuronal nuclei (**c**, FC: CTRL, 493 neurons, n = 4 vs PD, 418 neurons, n = 4; **d**, EC: CTRL, 224 neurons, n = 4 vs PD, 373 neurons, n = 4; **e**, CC: CTRL, 233 neurons, n = 4 vs PD, 247 neurons, n = 4). Data in graphs are reported as mean ± standard deviation. Mann-Whitney test, *p < 0.05. (**f, g**) PAR distribution in *substantia nigra* of control subjects and PD patients. The graph in **h** shows the percentage of PAR positive neuronal nuclei in *substantia nigra* (CTRL, 120 neurons, n = 4 vs PD, 150 neurons, n = 5). Data are reported as mean ± standard deviation. Mann-Whitney test, ns. (**i-l”**) show PAR and α-synuclein distribution in *substantia nigra* of control and PD patients. In control subjects, neuronal PAR staining is restricted to the nuclei (**i-i’**), and α-synuclein presents a dot-like signal within the synaptic compartment (**i,i”**). On the contrary, in PD, both neurons with PAR positive nucleus (**l**, white arrow) and neurons with PAR negative nucleus (**l**, dashed rectangle) are observed; in both cases, PAR is clearly detectable also in the neuronal cellular body, indicating a redistribution from the nucleus. α-Synuclein begins to accumulate in the neuronal cell body, colocalizing with PAR (**l”**). The graph in **m** shows the percentage of cells with PAR negative or positive nucleus (PAR^-^ and PAR^+^ nuclei, respectively) and with or without diffused or aggregated α-synuclein in the cytoplasm (α-syn^+^ and α-syn^-^ cytoplasm, respectively) of *substantia nigra* dopaminergic neurons (54 neurons, n=4 PD). (**n-o”**) show PAR distribution in neurons with undefined (n-n”) and ring-shaped (**o-o”**) α-synuclein aggregates. The graph in **p** shows the percentage of cells with PAR positive or negative nucleus presenting undefined (25 neurons, n=4 PD) or ring-shaped (19 neurons, n=4 PD) α-synuclein aggregates. Nuclei are counterstained using Hoechst. Scale bar, 20 μm. CC = cingulate cortex; CTRL = control; EC = entorhinal cortex; FC = frontal cortex; N = nucleus; ns = non-significant; SN = *substantia nigra*; white asterisks = neuromelanin.

We further investigated the relationship between PAR and α-synuclein in the *substantia nigra*. Thus, we conducted immunofluorescence for PAR, TH, and α-synuclein. As expected, in neurons of control subjects, PAR staining was confined to the nuclei, while we observed the classical α-synuclein dot-like signal primarily localized in the synaptic compartment (<u>Fig. 1i-i”</u>). In PD samples, we observed both neurons with PAR-positive nuclei (<u>Fig. 1l</u>, white arrow) and PAR-negative nuclei (<u>Fig. 1l</u>, dashed rectangle). In both cases, PAR was also clearly detectable in the neuronal cytoplasm, indicating its translocation from the nucleus (<u>Fig. 1l</u>). *a*-Synuclein began to accumulate in the cell body (<u>Fig. 1l, l”</u>), where PAR was also distributed (<u>Fig. 1l-l’</u>). Notably, among the neurons with PAR-positive nuclei, a significant proportion (76%) also exhibited diffuse or aggregated α-synuclein in the cytoplasm (<u>Fig. 1m</u>). To further explore these findings, we investigated PAR positivity in the nucleus alongside the presence of undefined (<u>Fig. 1n-n”</u>) or ring-shaped (<u>Fig. 1o-o”</u>) α-synuclein aggregates, which represents different stages of aggregate maturation 30,^31^. Interestingly, the percentage of neurons containing ring-shaped α-synuclein aggregates was higher in presence of PAR-positive nuclei (95%) compared to PAR-negative nuclei (56%) (<u>Fig. 1p</u>). Collectively, these data indicate that the redistribution of PAR from the nucleus to the cytoplasm occurs exclusively in neurons and appears to be associated to α-synuclein aggregation. Moreover, a second step of redistribution was observed, in which PAR distributes again in the nucleus with the appearance of ring-shaped α-synuclein aggregates in the cytoplasm. These data provide additional insights into the crucial steps of LB formation, advancing our understanding from the earlier stages involving undefined aggregates to the development of the characteristic ring-shaped structure^31–33^.

### 2.2 PARylation and stress granules co-work in α-synuclein aggregation during LB formation

We investigated in more detail the interplay between α-synuclein and PAR in undefined and ring-shaped aggregates, in dopaminergic neurons of substantia nigra (<u>Fig. 2 a-b”</u>). Interestingly, we observed that the fraction of α-synuclein that colocalizes with PAR significantly decreases from the undefined aggregate (<u>Fig. 2a-a”, c</u>), when the inclusion starts to form in the neuronal cytoplasm, to the ring-shaped aggregate, once α-synuclein is compacted to form a ring-like structure (<u>Fig. 2b-b”, c</u>).

**Fig. 2.**
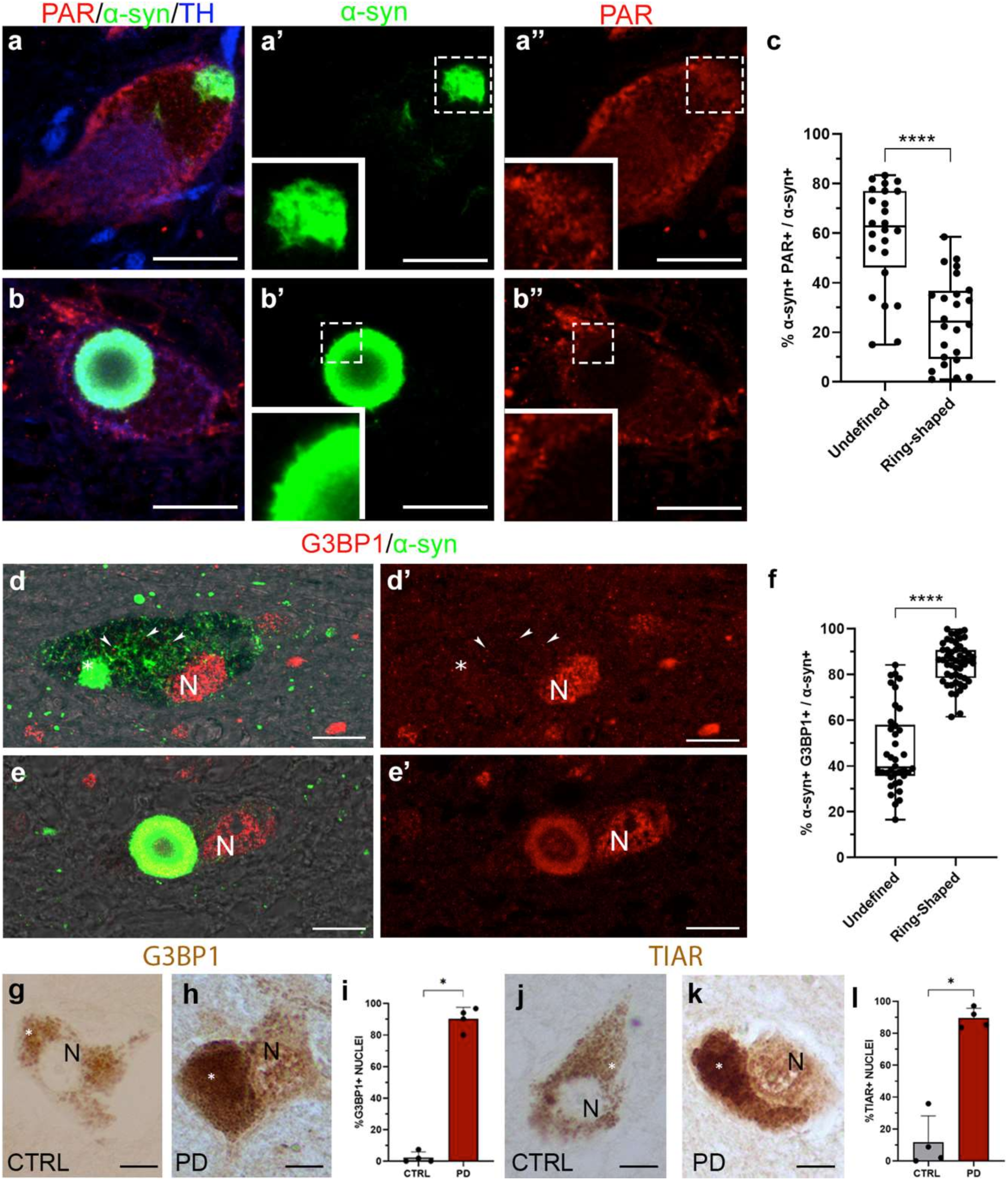
PAR and stress granules mutual involvement in α-synuclein aggregation in PD patients. (**a, b**) PAR translocate into the cell bodies of *substantia nigra* dopaminergic neurons and partially colocalize with α-synuclein aggregates. Colocalization is higher in the undefined stage (**a-a”**) compared with the ring-shaped structures (**b-b”**), where PAR localize in the peripheral region (**b”**). Scale bar, 20 μm. The graph in **c** (24 undefined aggregates, n = 4 PD; 24 ring-shaped aggregates, n= 4 PD) indicates the percentage of α-synuclein colocalizing with PAR expressed by Mander’s coefficient (M1). Data are reported as mean ± standard deviation. Mann-Whitney test, **** *p* < 0.0001. (**d** and **e**) show colocalization between α-synuclein and G3BP1, a marker of early stress granules. G3BP is present within the nucleus and shows a point-like signal in the cytoplasm (**d, d’**, arrowheads); stress granules are observed between the α-synuclein network (d’, arrowheads), evident in the cell body, and neuromelanin granules, visible with phase contrast. Colocalization increases from the undefined aggregate (**d, d’**, white asterisks) to the ring-shaped structure, where G3BP1 is located in the outer layer (**e-e’**). Scale bar, 20 μm. The graph in **f** (38 undefined aggregates, n = 5 PD; 51 ring-shaped aggregates, n = 5 PD) indicates the percentage of α-synuclein colocalizing with G3BP1 expressed by Mander’s coefficient (M1). Data are reported as mean ± standard deviation. Mann-Whitney test, **** *p* < 0.0001. (**g-l)** immunoenzymatic assay showing the distribution of G3BP1 and TIAR in *substantia nigra* dopaminergic neurons. In CTRL samples, nuclei are negative for both (**g, j**) while in PD patients, G3BP1 and TIAR localize within the nuclei (**h, k**). White asterisks indicate neuromelanin. Scale bar, 20 pm. The percentage of G3BP1 (CTRL, n = 4, 268 neurons vs PD, n = 4, 285 neurons) and TIAR (CTRL, n = 4, 229 neurons vs PD, n = 4, 274 neurons) positive nuclei are shown in **i** and **l**, respectively. Data in graphs are reported as mean ± standard deviation. Mann-Whitney test, *p <0.05; CTRL = control; N = Nucleus. Inset in a’, a”, b’, b”: 2,5x.

Since G3BP1 PARylation in the cytoplasm triggers stress granule assembly in cell models^15,18,19^, that can act as seed for pathological aggregation^34,35^, we investigated the presence of two key stress granule components, the RNA binding protein G3BP1 and TIAR, in LB formation. Analysing the distribution of G3BP1 and TIAR, the typical staining of small, roundish granules was observed in the neuronal cell bodies (<u>Fig. 2d’</u>, white arrowheads; Supplementary Fig. 2 a’). G3BP1 and TIAR colocalize with α-synuclein aggregates in both undefined aggregates (<u>Fig. 2d, d’,</u> Supplementary Fig. 2a, a’) and in the ring-shaped structure (<u>Fig. 2e, e’,</u> Supplementary Fig. 2b, b’). In contrast to PAR, quantitative analyses show a significant increase of G3BP1 in mature aggregates compared to undefined aggregates (<u>Fig. 2f</u>). We also observed an increase, although non-significant, for TIAR (Supplementary Fig. 2c). Moreover, we revealed a significant increase in the nuclear staining for both G3BP1 and TIAR in PD patients compared to controls (<u>Fig. 2g-l</u>).

All together these data indicate that PARylation and enrichment in stress granules are present in PD dopaminergic neurons and differently mark α-synuclein aggregates during LB formation.

### 2.3 PARylation involvement in early α-synuclein aggregation

Given the higher colocalization of PAR and α-synuclein in undefined α-synuclein aggregates (<u>Fig. 2c</u>), we hypothesized that PAR may play a key role in the early stages of protein aggregation. To investigate this, we explored the interplay between PARylation and α-synuclein oligomers, which are considered the earliest form of α-synuclein aggregates^29,36,37^. We used Proximity Ligation Assay (PLA) to detect α-synuclein oligomers alongside immunofluorescence in *substantia nigra* of PD patients. As expected, PAR was localised in the cell bodies of TH-positive neurons (<u>Fig. 3a, a’, b, b’, c, c’</u>), where α-synuclein oligomers were also present (<u>Fig. 3a, a’, b, b’, c, c’</u>). In detail, these oligomers, found throughout the cytoplasm in early aggregation stages (<u>Fig. 3a’</u>), as well as more aggregated forms (<u>Fig. 3b’</u>), and ring-shaped structure (<u>Fig. 3c’</u>), colocalized with cytoplasmatic PAR.

**Fig. 3.**
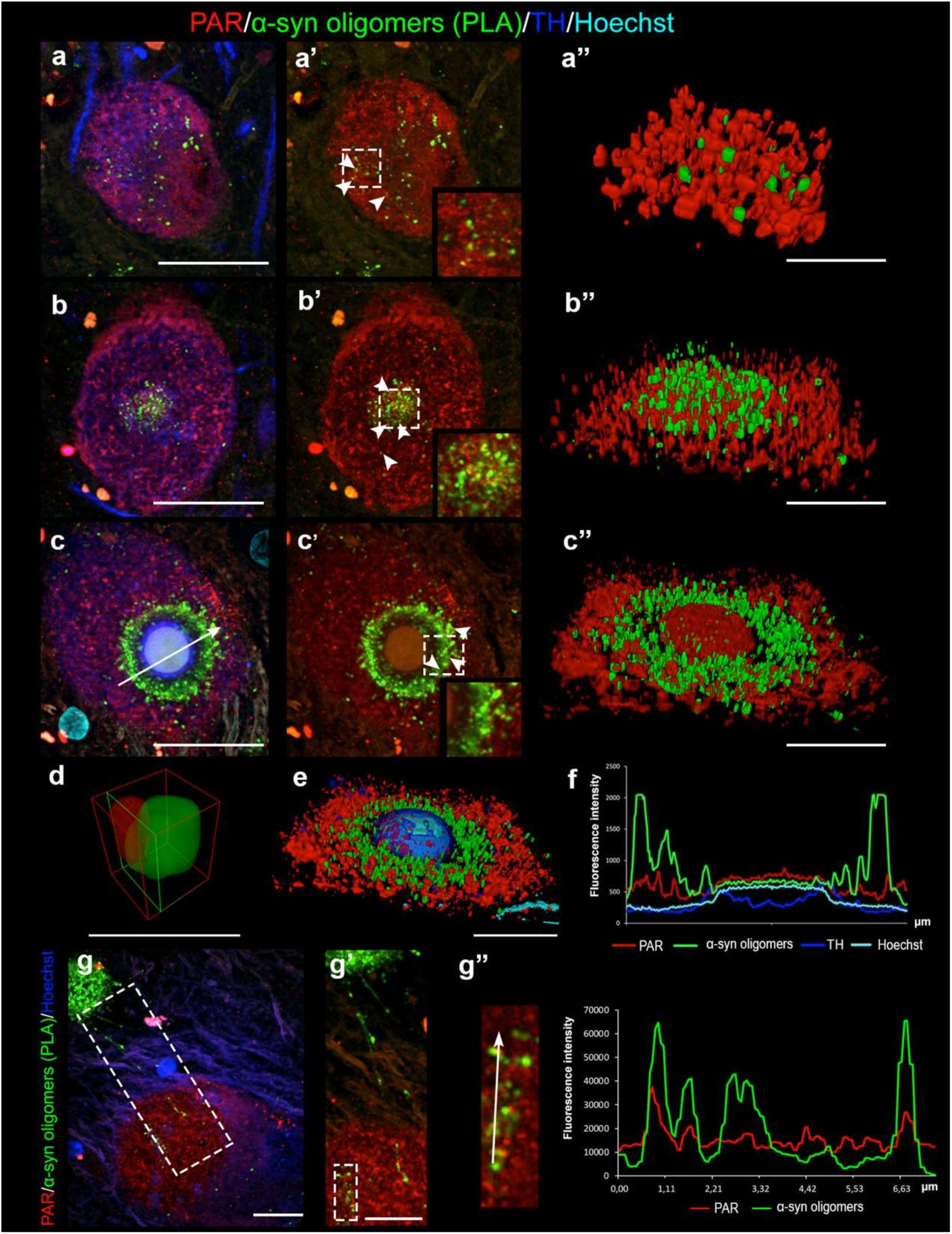
PAR colocalizes with α-synuclein oligomers during the aggregation process in PD patients. In *substantia nigra* dopaminergic neurons (**a-c’**), colocalization between PAR and α-synuclein oligomers in neuronal cell bodies, starting from the earliest stage, when α-synuclein oligomers are scattered in the cytoplasm (**a’**, white arrowheads), but also in undefined aggregate (**b’**, white arrowheads), and in the ring-shaped structure where colocalization is limited at the peripheral region (**c’**, white arrowheads). (**a”**, **b”** and **c”**) show 3D reconstruction obtained with arivis Vision4D 3.6.0 software of confocal images. (**d**) Example of PARylation of α-synuclein oligomer obtained with 3D reconstruction. (**e-f**) 3D reconstruction by arivis Vision4D 3.6.0 of **c** and the intensity profile graph for PAR, α-synuclein oligomer, TH and Hoechst signals along the line intersecting the ring-shaped aggregate in **c**. Nuclei are counterstained using Hoechst. (**g-g”**) α-synuclein oligomers outline a structure resembling a tunnelling nanotube (white dashed rectangle, 1,5x magnified in **g’**), which connects a cytoplasmic PAR-rich neuron and an α-synuclein aggregate. **g’’** shows the 2.5x magnification of a small portion of the tunnelling nanotube in **g’** (white dashed rectangle) and the intensity profile that shows partial overlapping peaks, indicating that also in this structure some oligomers are PARylated. **a, a’**, **b, b’**, **c**, **c’**: scale bar, 20 μm. **a’’**, **b’’**, **c’’**, **e**: scale bar, 5 μm. **d**: scale bar, 1 μm. **g**, **g’**: scale bar, 10 μm. Inset in a’, b’, c’: 1,5x.

To further analyze the interaction between PARylation and α-synuclein oligomers, we used high-resolution imaging and 3D reconstruction, and quantified α-synuclein oligomers and PAR interactions (<u>Fig. 3a”, b”, c”</u>, Supplementary video 1-3). Through this approach, we confirmed that α-synuclein oligomers colocalize with PAR (<u>Fig. 3d</u>) and quantified the percentage of PAR-positive oligomers. The percentage increased during the early and intermediate stages of aggregation (32% and 50% respectively) and decreased when ring-shaped structures formed (15%). Notably, the ring-shaped structure showed an onion-like structure with PAR distributed both in the center, where they colocalize with Hoechst, and in the periphery, where they colocalize with α-synuclein oligomers, surrounded by TH signal (<u>Fig. 3 e, f</u>). Moreover, a detailed examination of brain sections revealed the presence of some α-synuclein oligomers-positive structures (<u>Fig. 3g, g’</u>; n=2), that resemble the so-called tunnelling nanotube, recently described in human brain^31^. In detail, the neuron to which the tunnelling nanotube connects, is decorated with cytoplasmic PAR staining that colocalize with α-synuclein oligomers (<u>Fig. 3g”</u>), indicating a possible involvement of PARylation in the spreading of oligomers.

Finally, we examined the relationship between stress granule proteins G3BP1, TIAR, and α-synuclein oligomers, revealing that G3BP1 and TIAR-positive stress granules accumulate alongside α synuclein oligomers (Supplementary Fig. 3).

Collectively, these data support a pivotal involvement of PARylation in the earliest stages of α-synuclein aggregation.

### 2.4 PARylation is involved in Parthanatos and α-synuclein spreading in *substantia nigra* of PD patients

PAR accumulation in the cell body has been previously described as indicative of severe cellular stress^38^. Indeed, AIF PARylation induces its release from the mitochondria and, consequently, the activation of Parthanatos leading to cell death^23^. Evidence for the involvement of Parthanatos pathway in PD pathology has primarily been derived from *in vitro* studies and mouse models^6,39^. Therefore, in this work, we aimed to investigate whether PAR could localize into mitochondria in human PD brains and could be responsible for neuronal cell death of *substantia nigra* dopaminergic neurons. Triple immunofluorescence analysis revealed that mitochondria, marked by Voltage-dependent anion channel 1 (VDAC1), are widely distributed throughout the cytoplasm of dopaminergic neurons in both control (<u>Fig. 4a, a’</u>) and PD samples (<u>Fig. 4b, b’</u>). In control neurons, PAR predominantly localizes to the nucleus (<u>Fig. 4a, a’’</u>). However, in PD neurons, PAR accumulation to the cytoplasm (<u>Fig. 4b, b’’</u>), as previously demonstrated, and colocalizes with VDAC1 (<u>Fig. 4b</u>), as further confirmed by high-resolution 3D imaging (<u>Fig. 4e</u>). Interestingly, we observed that PAR also colocalize with mitochondria surrounding α-synuclein-labelled LB (<u>Fig. 4c-c’’</u>).

**Fig. 4.**
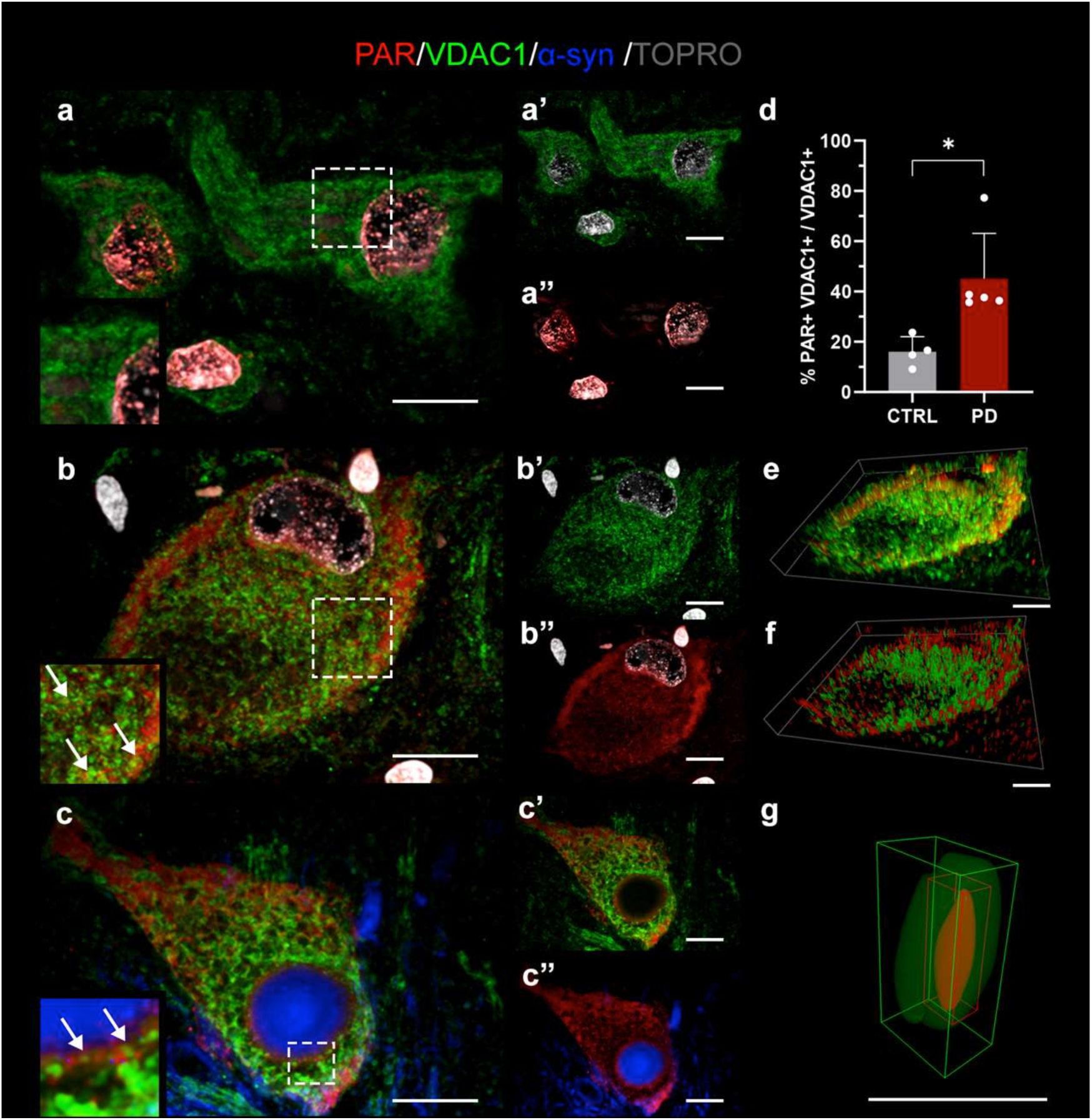
Mitochondria PARylation in *substantia nigra* dopaminergic neurons of PD patients. In control subjects, PAR are distributed in neurons nuclei (**a, a”**) while VDACl-labeled mitochondria localize into the cytoplasm, as expected (**a, a’**). In PD samples, cytoplasmic PAR (**b, b”**) partially colocalize with mitochondria (**b,** white arrows in the inset). PAR also colocalize with mitochondria that surround α-synuclein labeled LB (white arrows in the inset of **c,** 3x magnified). The graph in **d** (CTRL, n= 4, 88 neurons vs PD, n= 5, 133 neurons) indicates the percentage of mitochondria staining colocalizing with PAR expressed by Mander’s coefficient (M1). Data are reported as mean ± standard deviation. Mann-Whitney test, \**p* < 0.05. **e** and **f** show the 3D reconstruction obtained with Nis Elements software and with arivis Vision4D 3.6.0 software, respectively, of image **b**. (**g**) 3D reconstruction of a mitochondrion of *substantia nigra* dopaminergic neuron showing PAR inside it. Nuclei are counterstained using TOPRO**. a-a’’**, **b-b’’**, **c-c’’**, **e, f**: scale bar, 20 μm. **g**: scale bar, 3 μm.

Quantitative analysis showed a significant increase in the colocalization between PAR and mitochondria in the cytoplasm of *substantia nigra* dopaminergic neurons in PD patients compared to control (<u>Fig. 4d</u>). Moreover, 3D reconstruction provided the first evidence of PAR localization within mitochondria (<u>Fig. 4f, g</u>), suggesting its involvement in activating the Parthanatos cell death pathway in human PD brain.

Since Parthanatos pathway is known to trigger DNA condensation^22,23^, we further investigated this phenomenon. We revealed that condensed DNA was present inside *substantia nigra* neurons and that the condensed DNA is also PAR positive and localized in neurons showing mature LB (<u>Fig. 5a-a”’</u>). All together, these findings suggest that PAR plays a critical role in mediating neuronal cell death in substantia nigra dopaminergic neurons in PD patients.

**Fig. 5.**
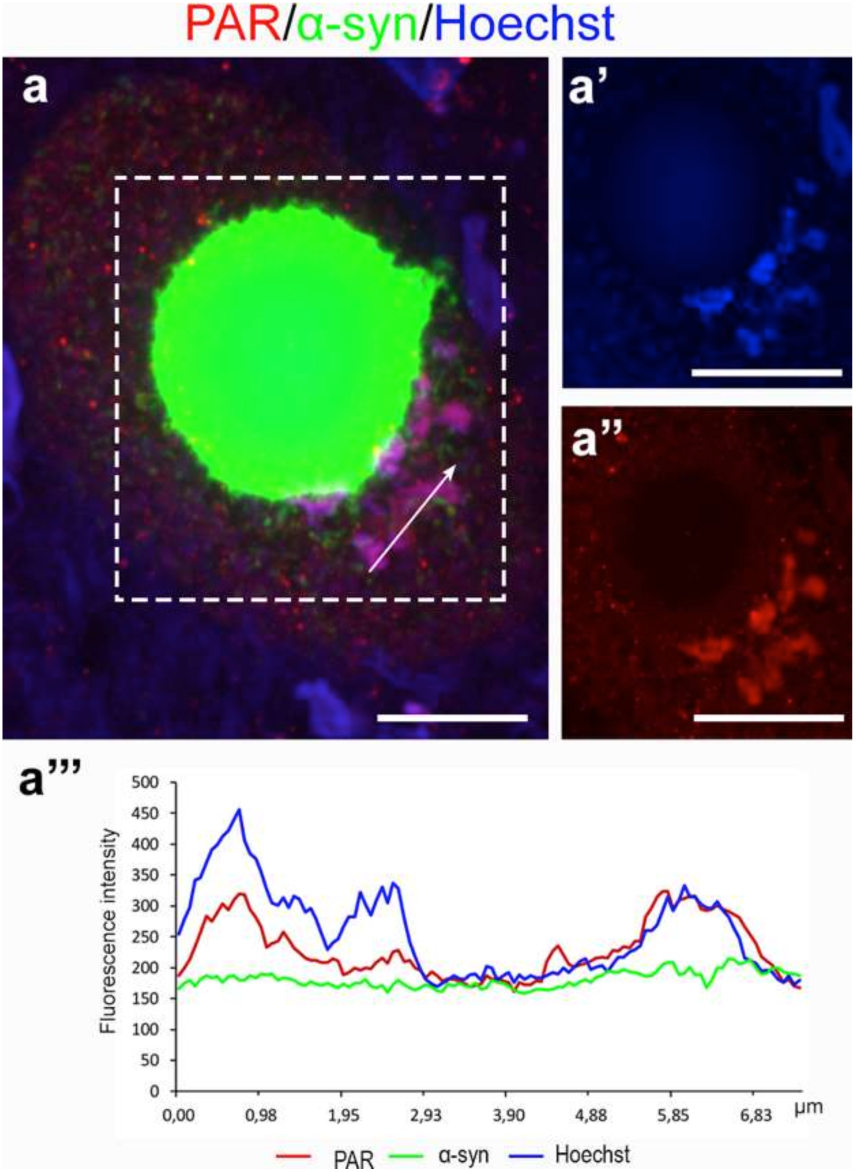
Interplay among α-synuclein aggregates, DNA condensation, and PARylation. *Substantia nigra* dopaminergic neuron with a mature LB (**a**) shows condensed DNA, marked with Hoechst (**a’**), that colocalizes with PAR (**a”**). (**a”’**) Intensity profile graph that highlights the overlapping signals of Hoechst and PAR along the arrow shown in **a**. Scale bar, 10 μm.

## 3 Discussion

PARP1-mediated PARylation has emerged as an important player in PD pathology, and, particularly, in exacerbating α-synuclein aggregation^6,28^. This key finding is supported by studies of cell and mouse models as well as in the CSF of PD patients^6,25^. Building on this, we investigated PARylation in *post-mortem* human PD brain to explore its potential link with α-synuclein aggregation and neuronal cell death. Our results led us to the proposed model illustrated in (<u>Fig. 6</u>). Briefly, in PD PAR polymers usually found in the nucleus (<u>Fig. 6a</u>) accumulate into the cytoplasm of neurons (<u>Fig. 6b</u>), where they colocalize with different forms of α-synuclein aggregates (<u>Fig. 6c-f</u>), mainly with the earliest form (<u>Fig. 6c</u>), and also in the periphery of LB (<u>Fig. 6d, e</u>) suggesting that PARylation may act as a trigger for α-synuclein aggregation. α-Synuclein aggregates colocalize also with stress granule proteins, though this occurs after the aggregates become PAR-positive (<u>Fig. 6c-e</u>), suggesting a role of PARylation in stress granules formation in PD human brain, as previously demonstrated in cellular and animal models^40,41^. Finally, PAR polymers colocalize with mitochondria having the potential to activate Parthanatos cell death pathway (<u>Fig. 6e</u>), as strongly suggested by the presence of condensed PARylated DNA (<u>Fig. 6f</u>). Studying *post-mortem* human PD brains provides valuable insight into the molecular events driving LB formation, complementing the knowledge gained from *in vitro* and animal models^30^. Our findings indicate a significant role for PAR in the neurodegenerative process of PD. This advances our understanding of the molecular events underlying LB formation, their propagation, and the mechanisms leading to neuronal cell death in PD patients.

**Fig. 6.**
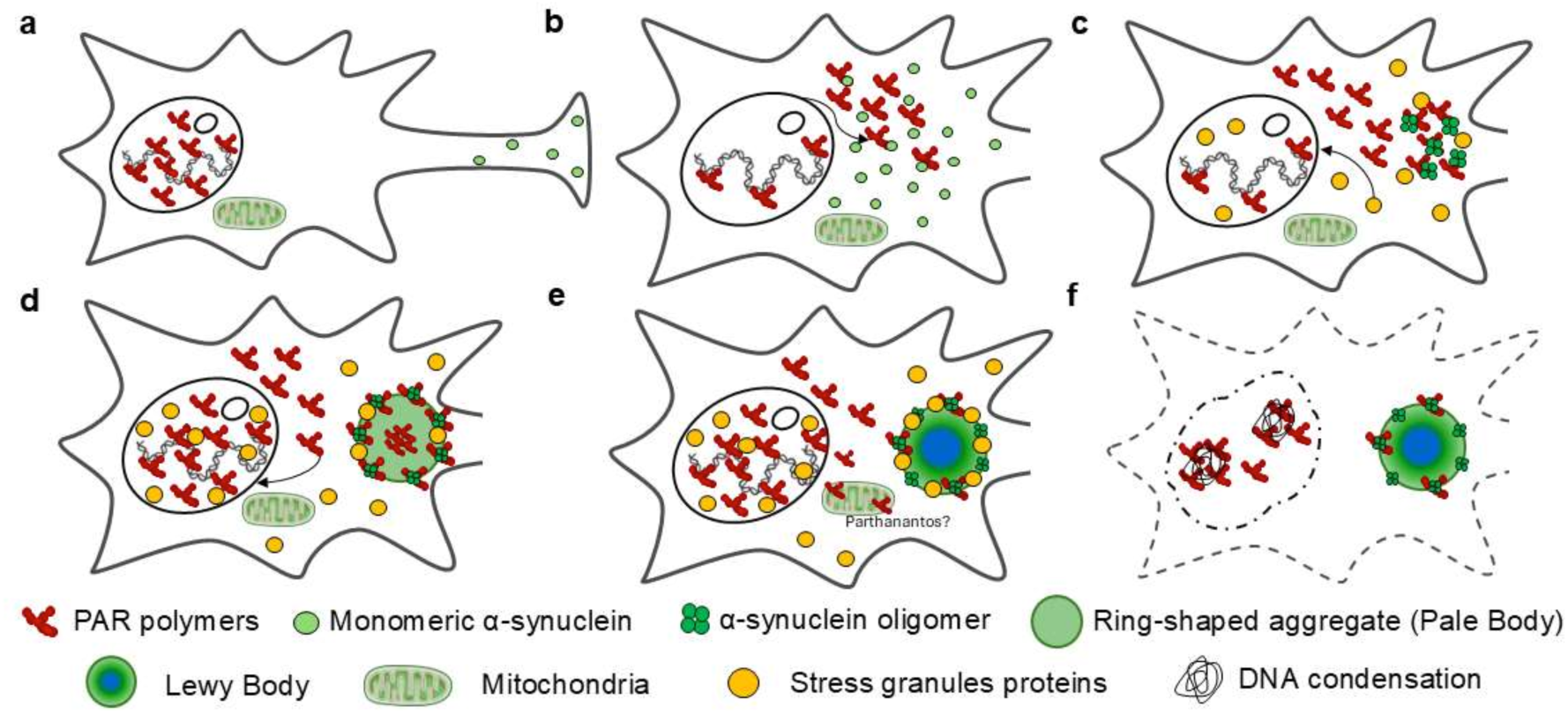
PAR redistribution and α-synuclein aggregation during Lewy body formation and neuronal cell death. Schematic model showing the sequences of events in LB formation concurrently with the redistribution of PAR and stress granules. (**a**) In controls, PAR are located in the nuclei, where they carry out their physiological function in repairing DNA, while α-synuclein mainly localizes in the synaptic compartment. (**b**) In PD, both PAR and α-synuclein start to accumulate in the neuronal cell body. (**c**) α-synuclein oligomers are present in undefined aggregates where colocalize with PAR and stress granules proteins (Supplementary Fig. 3). Stress granules partially translocate within the nucleus. (**d**) With the formation of aggregates with ring-shaped structure, both PAR and stress granule proteins localize within the nucleus; notably PAR are arranged both in the center and in the periphery of the ring-shaped structure (Pale Body), where they co-localize with α-synuclein oligomers. (**e**) PAR polymers co-localize with mitochondria, which is the first step in activating the Parthanatos cell death pathway; in the mature LB, PAR and stress granule proteins can be identified in the peripheral region; Hoechst staining (blue) is detectable in the core of these aggregates. (**f**) Neurons that show a mature LB and, also show PARylated and condensed DNA.

Although alterations of PARPl-mediated PARylation have been extensively described in many pathological conditions such as cancer, autoimmune, and cardiovascular diseases^13^, its role in neurodegenerative diseases including PD, Alzheimer’s disease, Huntington’s disease, and multiple sclerosis is emerging only in these last years^42^. PARP1 is primarily localized in the nucleus, where it plays a protective role in repairing DNA single and double strand breaks. However, prolonged activation of PARP1 can lead to cell death through NAD+ depletion, impaired cellular function, and DNA fragmentation^7^. Thus, PARP1 overactivation, observed in PD, suggests that PAR overproduction may be a key pathological process^13,15^. In this study, we observed a decrease of PAR in the neuronal nuclei of PD patients compared to control in cortical areas affected by α-synuclein pathology and a strong translocation from the nucleus to neuronal cytoplasm, in both the cortex and *substantia nigra*. This cytoplasmic redistribution aligns with the nuclear depletion and cytoplasmic relocation of the PARP1 previously reported in dopaminergic neurons from *post-mortem* PD brains^27^ as well as the increased PAR levels detected in the CSF of PD patients^6^.

In this study, we found that PAR in the soma of *substantia nigra* dopaminergic neurons colocalize with cytoplasmic α-synuclein and with the different forms of α-synuclein inclusions during LB morphogenesis. Over the years, many efforts have been made to identify the different components of LB and understand the mechanisms leading to their formation^30,31^. Furthermore, the higher colocalization of PAR with the early stages of α-synuclein aggregation, together with the presence of PAR-positive α-synuclein oligomers, highlights a potential role for PARylation in triggering α-synuclein aggregation, and supports prior hypotheses that PARP1-mediated PARylation is involved in this process. PARylation may drive aggregation through direct interaction between PAR and α-synuclein, likely via electrostatic forces involving the positively charged lysine residues of α-synuclein 28. Alternatively, PARylation may indirectly affect α-synuclein aggregation through its regulation of autophagy which is impaired in PD^43^. Overactivation of PARP1 and an overproduction of PAR, as reported in the present work, could result in PARylation of autophagy-related proteins and NAD+ depletion that induce SIRT1 inactivation^44,45^. This could impair autophagy and increase the susceptibility of neurons to aggregated α-synuclein, damaged organelles, and oxidative stress. Additionally, PAR elevation in the cytoplasm has been shown to promote stress granule formation in cell models^18^, with PARylation promoting granule biogenesis^19^. For the first time, we observed the presence of stress granule proteins in *substantia nigra* dopaminergic neurons of PD human brain, particularly in mature LB and, to a lesser extent, in undefined aggregates. The higher presence of stress granules in LB compared to earlier forms, suggest that stress granules assembly may occur later, while PAR, plays a more pivotal role in the early steps of PD pathology.

G3BP1 ADP-ribosylation has previously been described in cell models as a key actor in stress granules formation^18^ under both acute and chronic stress conditions^46^, and is also implicated in the nuclear translocation of G3BP1^19^. In our study, alongside stress granule formation, we observed the increase in the nuclear staining for both G3BP1 and TIAR in PD patients compared to controls, indicating a nucleocytoplasmic shuttling for these two proteins. This phenomenon may reflect the chronic stress characteristics of PD. Indeed, the role of PARylation in the nucleocytoplasmic shuttling has been already demonstrated for another stress granule protein, the adenylateuridylate-rich element (ARE)-binding protein embryonic lethal abnormal vision-like 1 (Elavl1)/human antigen R (HuR), a mechanism to regulate gene expression at the post-transcriptional level in an *in vitro* model^47^. The observed nuclear-cytoplasmic dynamics of G3BP1 and TIAR in PD may, therefore, indicate a broader role for PARylation in modulating cellular stress responses and gene expression. This supports the hypothesis that chronic PARP1 activation in PD may contribute to the pathological aggregation of α-synuclein and stress granule formation through these nucleocytoplasmic transport mechanisms.

The role of α-synuclein neurodegeneration in PD via the Parthanatos death cascade was first described following *in vivo* and *in vitro* administration of α-synuclein pre-formed fibrils (PFFs) that increases nitric oxide formation and activates PARP1^6^. This was later confirmed by studies showing that genetic and pharmacological manipulation of the pathway prevents neurodegeneration in PD mouse models and primary neuronal cultures^39^. In *post-mortem* human brain, we observed a marked increase in PAR colocalization with mitochondria in *substantia nigra* dopaminergic neurons of PD patients compared to controls. Moreover, through 3D reconstruction we identified PAR polymers within mitochondria.

Given that endogenous AIF, one of the effectors of Parthanatos pathway, is located both at the outer mitochondrial membrane on the cytosolic side and attached to the inner membrane facing the intermembrane space 48, and binds PAR with high affinity 49, our findings suggest that the increased mitochondria PARylation in dopaminergic neurons of PD patients, may indicate Parthanatos pathway activation. Additionally, we also observed PAR colocalization with mitochondria adjacent to a mature LB. This might indicate that a link exists between α- synuclein, LBs, and neuronal cell death. Neurons with mature LBs also displayed PARylated condensed DNA, further supporting this connection.

In light of these results and emerging data from the literature, it is clear that PARylation could represent an attractive therapeutic target to act on in PD pathology^38^. Two main approaches could be pursued: *i)* the direct inhibition of PARP1 and *ii)* the inhibition of PARP1 effectors in the Parthanatos pathway. The first approach could be based on the inhibition of PARP1 enzyme to counteract the production of excessive PAR polymers that could exacerbate α-synuclein aggregation^25^. Several inhibitors are commercially available and currently used to treat different types of cancer, acting as a PARP1 active site binder or as trapping compound on DNA. Among them, olaparib and veliparib, used in the treatment of ovarian and breast cancer, were also found to be protective in different *in vitro* and *in vivo* PD and neurodegenerative models^25,50^. Moreover, administration of PARP1 indirect inhibitors (nilotinib) has been attempted in a clinical trial, in PD patients, but did not provide the expected results^51^ (ClinicalTrial.gov: NCT03205488). This is understandable since patients enrolled were in the late stage of the disease, when PARP1 has already exerted its action and, probably, contributed to α-synuclein aggregation and neuronal cell death via Parthanatos pathway. Therefore, it might be an option to consider PARP1 as an early target to act on in the early stages of neurodegeneration. Nevertheless, it cannot be excluded that lack of effect is due to the inhibitor used that may not acts directly on PARP1. However, some issues need to be considered; among these, the selectivity is fundamental, since some of the inhibitors act not only on PARP1 but also on PARP2 and other PARP isoforms. Moreover, the administration could have to be chronic, and, given PARP1 plays many biological functions, this may be associated with a risk of various side effects such as genotoxicity^50^. Recently, another compound, called 10e, was tested *in vitro* and emerged as a potential inhibitor to be further investigated in PD animal models. Indeed, it is a structural analogue of FDA-approved olaparib, with higher PARP1 affinity, selectivity, and lower toxicity^52^. Since it appears that the death of *substantia nigra* dopaminergic neurons is mediated by the PARP1-based cell death called Parthanatos pathway, the second approach that could be pursued is to inhibit the effectors of this pathway such as the flavoprotein AIF and the nuclease MIF. Thus, the strategies of controlling the level of intracellular PAR targeting Poly (ADP-ribose) glycohydrolase (PARG) enzyme, which is responsible of its degradation^24^, might be considered.

In conclusion, this study points out the role of PARP1-mediated PARylation in PD pathology. The study of molecular events occurring in *post-mortem* human brain integrated with the experimental studies on cell and mouse models, indicate a role of PAR in stress granule assembly and LB formation, their propagation and, ultimately neuronal cell death. The involvement of PARylation in triggering α-synuclein aggregation, which is supported by the identification of PARylated α-synuclein oligomers, and in triggering Parthanatos cell death pathway in *substantia nigra* dopaminergic neurons, suggests that it could be a therapeutic target in an early stage on the disease. Finally, a question arises: what can cause chronic stress, inducing PAR overactivation and stress granules formation, as observed in PD patients? Some studies reported a possible involvement of viral infection in PD^53^ and viral infectious has been demonstrated to be able to activate stress granules^54,55^.

## 4 Methods

### 4.1 Patients

All patients were enrolled and followed during the course of the disease at the Parkinson’s Centre ASST G. Pini-CTO of Milan. The clinical diagnosis of PD was carried out according to the UK Brain Bank criteria^56,57^ and confirmed by neuropathological assessment performed by two experienced neurologists in movement disorders, GG and MB, in compliance with current BrainNet Europe Consortium guidelines^58^.

*Post-mortem* human brains obtained from PD patients (N=8) and from age-matched control subjects (N= 6) clinically free from neurological diseases were used (Supplementary Table 1). Written informed consent was obtained from all subjects in agreement with relevant laws and institutional guidelines and approved by the appropriate institutional committees. The study procedures were approved by the Ethics Committee of the University of Milan (protocol code 66/21).

Brains were fixed in 10% buffered formalin for at least 21 days at 20°C. Selected areas were paraffine embedded and cut at the microtome (MR2258, Histoline) to obtain 5 mm thick sections of mesencephalon, entorhinal, cingulate and the frontal cortex, processed for the following analysis.

### 4.2 Immunohistochemistry

After deparaffination and rehydration, tissue sections were sequentially incubated with: *i)* 80% formic acid solution for 20 min for antigen retrieval, *ii)* 3% H2O2 for 20 min for endogenous peroxidases inactivation; *iii)* 1% BSA in 0.01 M phosphate saline buffer (PBS) containing 0.1% Triton X-100 for 20 min (BSAT); *iv)* the primary antibody, mouse anti-PAR (the PAR chain length used as immunogen ranges from 2 to 50 monomers; 1:200), mouse anti-G3BP (1:250) or mouse anti-TIAR (1:100), diluted in BSAT and incubated overnight at room temperature (RT) (Supplementary Table 2). To visualize the antigen-antibody binding the EnVision anti-mouse secondary antibody (Agilent 1 h, RT) and 3,3‘-Diaminobenzidine as chromogen (DAB, Agilent kit) were used (Supplementary Table 2). Tissue sections were mounted with permanent mounting medium (Eukitt). Images were acquired with the optical microscope Zeiss at 10X and 20X magnification.

### 4.3 Immunofluorescence and Proximity Ligation Assay (PLA)

For the immunofluorescence staining, sections containing *substantia nigra* and frontal cortex were incubated with either 80% formic acid solution for 20 min or high pH antigen retrieval solution 50X (Agilent, S2367; diluted 1:50 in PBS) at 90°C for 20 min plus 10 min at RT and 5 min in PBS. Then, incubation with BSAT for 20 min at RT followed by a mix of anti-PAR with different primary antibodies (S100 calcium-binding protein B (S100β), Tyrosine Hydroxylase (TH), Voltage-dependent anion channel 1 (VDAC1), and α-synuclein), or by a mix of G3BP1/TIAR and α-synuclein antibodies was carried out overnight at RT (Table 2). Tissue sections were then incubated with specific fluorescent secondary antibodies (Table 2) for 2 h at RT in the dark.

PLA was used to detect α-synuclein oligomers^36^, accordingly to protocols previously described^59^. Briefly, sections were incubated with a mixture containing α-synuclein S3062-MINUS and α-synuclein S3062-PLUS probes, anti-PAR and anti-TH antibody diluted in Duolink® Antibody Diluent for 1 h at 37°C and then overnight at RT. To carry out the amplification reaction, tissue sections were treated as follows: *i)* Duolink® ligase (1:40) in Duolink® ligation solution (1:5) (diluted in milliQ water at 37°C for 1 h, *ii)* Polymerase (1:80) in Duolink® Amplification Reagent Green (1:5 in milliQ water) for 2 h at 37 °C, to which donkey anti-mouse secondary antibody conjugated to Alexa Fluor® 568 and donkey anti-goat secondary antibody conjugated to Alexa Fluor® 647 were added to the polymerase step to detect double immunofluorescence. Nuclei were stained using either Hoechst 33342 (1:5000) or TO-PRO®-3 (1:1000), 10 min at RT. Sections were mounted using Mowiol® + DABCO® and finally, examined with Nikon spinning disk confocal microscope, equipped with CSI-W1 confocal scanner unit using a water-immersion 40X and silicon-immersion 100X objectives.

### 4.4 3D reconstruction

For 3D visualization, arivis Vision4D® 3.6.0 software (Zeiss Company), a powerful tool that guarantees precise and reproducible quantitative morphometric analyses to visualize small and complex structure^60,61^, was used. In detail, images were imported into the software and transformed into the 12-pixel format. The region of interest (ROI) of deconvolved 100X acquisitions was created using the “Transformation gallery > Crop” tool. After gamma correction to enhance faint objects and denoising to remove residual noise, different pipelines were used to 3D reconstruction. The “Intensity threshold segmentation” pipeline was used for both TH and nuclei continuous staining. PLA puncta, PAR and VDAC1 staining, which show a point like distribution, are reconstructed using “blob finder” pipeline, selecting a suitable range of exposure using the “preview” tool.^35,36^

### 4.5 Image and statistical analysis

Images were analyzed using Fiji software (NIH). In detail, counting of PAR positive nuclei of neurons and astrocytes in the corresponding anatomic regions was performed using cell counter tool, while JACOP plug-in was selected for colocalization analysis calculating Mander’s coefficient^62^. Statistical analysis was performed using GraphPad^TM^ 8.0 software. Un-paired non-parametric Mann-Whitney test was carried out to compare groups considering a *p-value* <0.05 as statistically significant.

## ACKNOWLEDGEMENTS

The authors thank all patients and families for their contribution, “Fondazione Pezzoli per la Malattia di Parkinson” (Milan-Italy) for SM, FG and AMC salary and for long-lasting support to GC.

## AUTHOR CONTRIBUTIONS STATEMENT

All authors contributed to the study conception. MB, EB, GG, SM, FG processed human samples. CN and FG performed immunohistochemistry and multiple labelling experiments. CN, SM and FG performed confocal and super resolution imaging, as well as processing and analysis of images, also using arivis Vision 4D software. SM, CN, FG, AMC, CR and GC performed data analysis. GC, GP, SM, CN, FG, GG, IUI, AMC, DC, EC and CR designed research, analysed and interpreted the data, and contributed to writing the manuscript.

## ETHICS STATEMENT

The study was conducted according to the guidelines of the Declaration of Helsinki and approved by the Ethics Committee of the University of Milan (protocol code 66/21,15 June 2021).

## DATA AVAILABILITY

The datasets generated and/or analyzed during the current study are available upon request from the corresponding authors.

## MATERIALS AND CORRESPONDENCE

Correspondence and requests for materials should be addressed to Graziella Cappelletti and Samanta Mazzetti

## SUPPLEMENTARY INFORMATION

### Supplementary Figures 1-3

Supplementary Figure 1: Confocal analysis of PAR in astrocytes.

Supplementary Figure 2. Confocal analyses of TIAR and α-synuclein during Lewy body morphogenesis in PD patients.

Supplementary Figure 3. Stress granules and α-synuclein oligomers.

### Supplementary Tables 1-2

Supplementary Table 1. Demographic and clinical characteristics of the subjects included in this study.

Supplementary Table 2. Primary and secondary antibodies and kits used in this study.

### Supplementary movies

Movie 1: The movie refers to arivis 4D software 3D reconstruction of figure 3a’’’.

Movie 2: The movie refers to arivis 4D software 3D reconstruction of figure 3b’’’.

Movie 3: The movie refers to arivis 4D software 3D reconstruction of figure 3c’’’.

