## Supplementary Figure 1-3; Supplementary Table 1-2 for "PARylation in Parkinson’s disease: a bridge between Lewy body formation and neuronal cell death"

#### Supplementary Figures 1-3

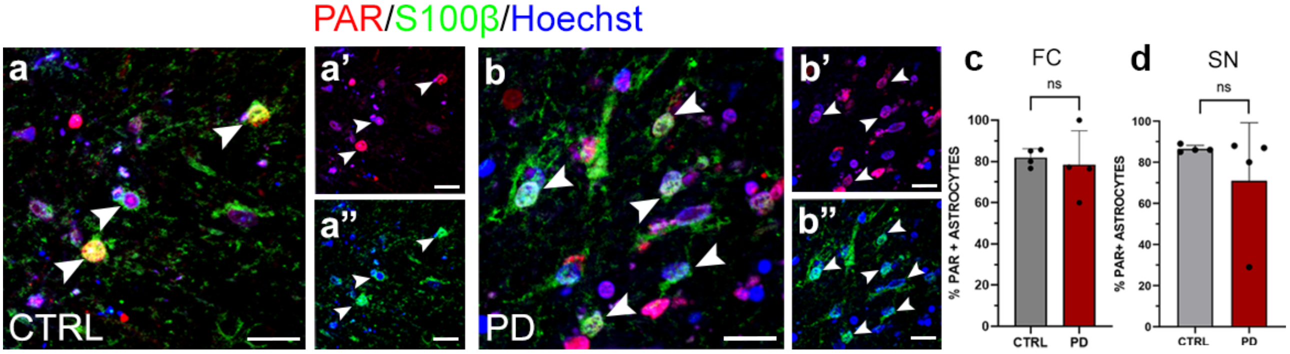


**Supplementary Figure 1.** Confocal analysis of PAR in astrocytes. (**a-b''**) representative image showing PAR nuclear distribution (white arrowheads) in astrocytes of *substantia nigra* of control subjects (**a-a''**) and PD patients (**b-b''**). Scale bar, 20 μm. The graphs show the percentage of PAR positive astrocytes nuclei in frontal cortex (**c**, FC: CTRL, 132 astrocytes, n = 4 vs PD, 171 astrocytes, n = 4) and *substantia nigra* (**d**, SN: CTRL, 141 astrocytes, n = 4 vs PD, 150 astrocytes n = 4). Data are reported as mean ± standard deviation. Mann-Whitney test, p>0.05; ns. CTRL = control; FC = frontal cortex; N = nucleus; ns = non-significant; SN = *substantia nigra*.

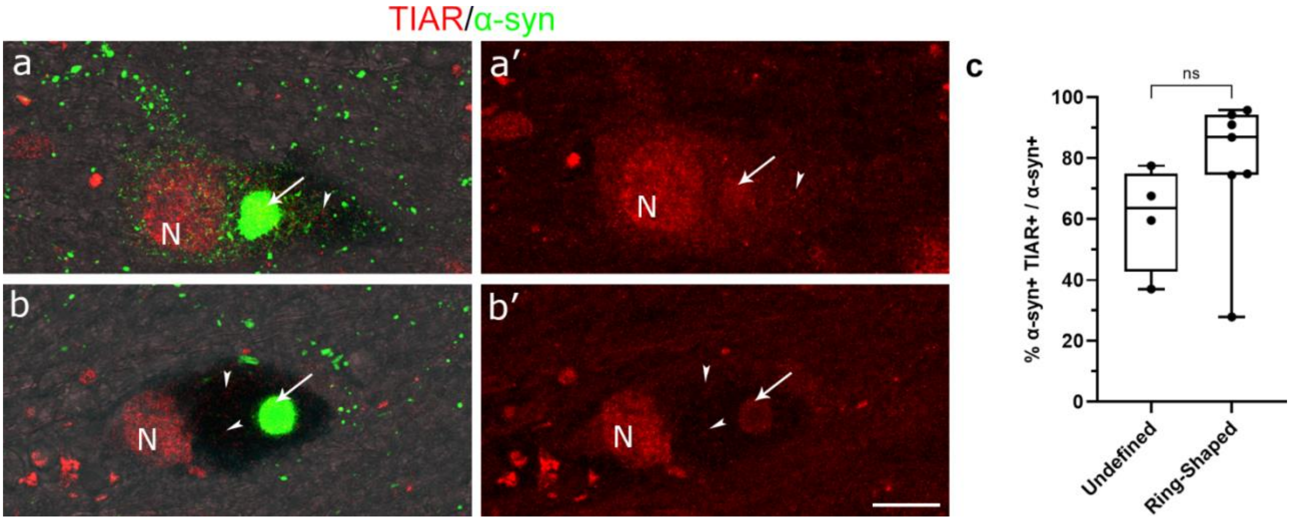


**Supplementary Figure 2.** Confocal analyses of TIAR and α-synuclein during Lewy body morphogenesis in PD patients. (**a, a'**) TIAR localizes in the nuclei and shows a point-like signal in the cytoplasm (white arrowhead) between the α-synuclein network and neuromelanin granules, visible with phase contrast, and colocalizing with α-synuclein undefined aggregate (white arrow). (**b, b'**) When the ring-shaped structure is formed TIAR is present in both the nucleus and cytoplasm (white arrowheads) where it colocalizes with α-synuclein (white arrow).

Colocalization increases from the undefined to the ring-shaped aggregate. Scale bar, 20 µm. The graph in **c** (n= 1 PD, undefined = 4; ring-shaped = 7) indicates the percentage of α-synuclein colocalizing with TIAR expressed by Mander's coefficient (M1). Data are reported as mean ± standard deviation. Mann-Whitney test, ns. N = Nucleus. ns = non-significant.

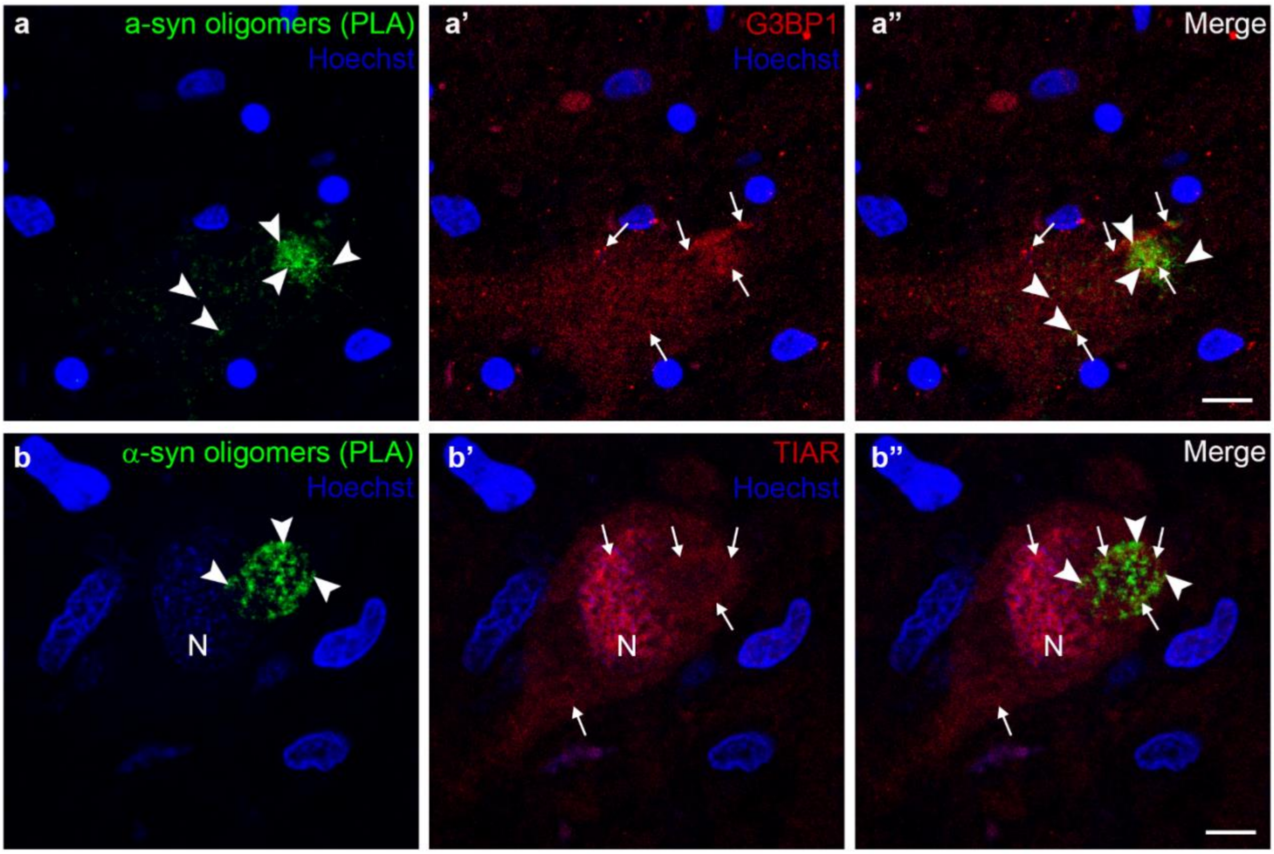


**Supplementary Figure 3.** Stress granules and α-synuclein oligomers. α-Synuclein oligomers are diffused in the cytoplasm and form an undefined aggregate (**a, b**; arrowheads); G3BP1 (**a'**) and TIAR (**b'**), showing a point-like signal (arrows) both in the cytoplasm and in the neuronal nucleus, partially colocalize with α-synuclein oligomers (**a", b"**). Nuclei are counterstained using Hoechst. Scale bar, 20 µm. N = Nucleus

### Supplementary Tables 1-2

| Clinical Diagnosis | Gender | Age of onset | Age of death | Disease duration (year) |
|---|---|---|---|---|
| CTRL#1 | M | / | 71 | / |
| CTRL#2 | F | / | 93 | / |
| CTRL#3 | F | / | 82 | / |
| CTRL#4 | F | / | 64 | / |
| CTRL#5 | F | / | 84 | / |
| CTRL#6 | F | / | 64 | / |
| PD#1 | M | 62 | 80 | 18 |
| PD#2 | M | 59 | 87 | 28 |
| PD#3 | F | 65 | 84 | 19 |
| PD#4 | F | 53 | 91 | 38 |
| PD#5 | F | 59 | 79 | 20 |
| PD#6 | M | 57 | 71 | 14 |
| PD#7 | M | 57 | 75 | 18 |
| PD#8 | M | 59 | 75 | 16 |

**Supplementary Table 1.** Demographic and clinical characteristics of the subjects included in this study.

| Antigen | Code | Host | Dilution |
|---|---|---|---|
| **Primary antibodies** | | | |
| G3BP (Human G3BP1 aa 200-350) | Ab56574 Abcam | Mouse | 1:250 |
| Poly-ADP ribose (PAR) | MABC547 Merck | Mouse | 1:100 |
| S-100β (Rat S100β aa 1 to 92) | 2874006 Synaptic Systems | Chicken | 1:500 |
| TIAR (Human TIAR aa 161-365) | 610352 BD Transduction Laboratories™ | Mouse | 1:100 |
| Tyrosine Hydroxylase (TH) (synthetic peptide sequence, VQDELDTLAHAL, corresponding to C-terminus) | PA518372 Thermo Fisher Scientific | Goat | 1:200 |
| VDAC1 (the immunogen used is proprietary information) | ab15895 Abcam | Rabbit | 1:200 |
| α-synuclein (human aa 111-132, corresponding to C-terminus) | S3062 Merck | Rabbit | 1:2000 |
| α-synuclein (human synthetic peptide aa 100 to C-terminus) | Ab21976 Abcam | Sheep | 1:100 |

| | | | |
|---|---|---|---|
| **Secondary antibodies** | | | |
| Alexa Fluor® 568 anti-mouse | A10037 Thermo Fisher Scientific | Donkey | 1:200 |
| Alexa Fluor® brilliant violet 421 anti-chicken | 703-675-155 Jackson Immunoresearch | Donkey | 1:300 |
| Alexa Fluor® 647 anti-goat | 705-615-147 Jackson Immunoresearch | Donkey | 1:300 |
| Alexa Fluor® 488 anti-rabbit | A21206 Thermo Fisher Scientific | Donkey | 1:200 |
| Alexa Fluor® 647 anti-rabbit | A32795 Thermo Fisher Scientific | Donkey | 1:200 |
| Alexa Fluor® brilliant violet 421 anti-sheep | 713-675-147 Jackson Immunoresearch | Donkey | 1:300 |
| **Probes, kit and stains** | | | |
| Duolink® in situ probe marker MINUS | DUO920101KT Merck | - | - |
| Duolink® in situ probe marker PLUS | DUO920091KT Merck | - | - |
| Duolink® In Situ Detection Reagents green | DUO92014 Merck | - | - |
| EnVision FLEX DAB + Substrate Chromogen System | K3468 Agilent | - | - |
| Hoescht 33342 | 62249 Thermo Fisher Scientific | - | 1:5000 |
| TO-PRO®-3 | 629661 Thermo Fisher Scientific | - | 1:1000 |

**Supplementary Table 2.** Primary and secondary antibodies and kits used in this study.
